# Discovery of therapeutic agents targeting *PKLR* for NAFLD using drug repositioning

**DOI:** 10.1101/2022.02.15.480557

**Authors:** Cheng Zhang, Mengnan Shi, Woonghee Kim, Muhammad Arif, Martina Klevstig, Xiangyu Li, Hong Yang, Cemil Bayram, Ismail Bolat, Özlem Özdemir Tozlu, Ahmet Hacımuftuoglu, Serkan Yıldırım, Yongjun Wei, Xiaojing Shi, Jens Nielsen, Hasan Turkez, Mathias Uhlen, Jan Boren, Adil Mardinoglu

## Abstract

**Background & Aims:** Non-alcoholic fatty liver disease (NAFLD) encompasses a wide spectrum of liver pathologies. However, not medical treatment has been approved for the treatment of the disease. In our previous study, we found *PKLR* could be a potential target for treatment of NALFD. Here, the aim is to investigate the effect of *PKLR* in *in vivo* model and perform drug repositioning to identify a drug candidate for treatment of NAFLD.

**Methods:** Biopsies from liver, muscle, white adipose tissue and heart were obtained from control and *PKLR* knockout mice fed with chow and high sucrose diets. Lipidomics as well as transcriptmics analyses were conducted using these tissue samples. In addition, a computational drug repositioning analysis was performed and drug candidates were identified. The drug candidates were finally tested in both *in vitro* and *in vivo* models to evaluated their toxicity and efficacy.

**Results:** The *Pklr* KO reversed the increased hepatic triglyceride level in mice fed with high sucrose diet and partly recovered the transcriptomic changes in liver as well as other three tissues. Both liver and white adipose tissues exhibited dysregulated circadian transcriptomic profiles, and these dysregulations were reversed by hepatic knockout of *Pklr*. In addition, 10 small molecule drugs were identified as potential inhibitor of *PKLR* by the drug repositioning pipeline, and two of them significantly inhibited both the *PKLR* expression and triglyceride level in *in vitro* model. Finally, the two selected small molecule drugs were evaluated in *in vivo* rat models and it was demonstrated that these drugs attenuated hepatic steatosis without side effect on other tissues.

**Conclusion:** In conclusion, our study provided biological insights about the critical role of *PKLR* in NAFLD progression and proposed a treatment strategy for NAFLD patients, which has been validated in preclinical experiment.

## Introduction

Non-alcoholic fatty liver disease (NAFLD) is defined as the accumulation of fat in the liver due to the imbalance in the uptake and secretion of fat as well as the increased *de novo* lipogenesis (DNL) or decreased oxidation of fat in the liver (Gluchowski et al., 2017). NAFLD can progress to non-alcoholic steatohepatitis (NASH) and it is a well-known risk factor for many metabolic diseases such as type-2 diabetes and cardiovascular disease (Mardinoglu et al., 2018; Tana et al., 2019). It has been reported that the prevalence of NAFLD is approximately 25% of the population, but it has no approved effective pharmacological treatment. Hence, there is an urgent need for the development of an effective treatment strategy.

Previously, we have generated gene co-expression networks (CNs) for the liver and other 45 primary human tissues, and identified the pyruvate kinase L/R (*PKLR*) as a target, whose inhibition may selectively inhibit DNL in the liver (Lee et al., 2017). Pyruvate kinase is a key enzyme in glycolysis and the *PKLR* gene encodes for the liver (*PKL*) and erythrocyte (*PKR*) isoforms of the enzyme and catalyses the production of pyruvate and ATP from phosphoenolpyruvate and ADP. *PKL* and *PKR* isoforms are specifically expressed in the liver and erythrocytes, respectively. An independent mouse population study has also verified the driving role of *PKLR* in the development of NAFLD (Chella Krishnan et al., 2018). Recently, we have performed *in vitro* experiments by inhibiting and overexpressing *PKLR* in HepG2 cells, and found that the expression of *PKLR* was significantly positively correlated with the expression of *FASN*, DNL, TAG levels and cell growth (Liu et al., 2019). In light of these findings, the integrative analysis suggested that *PKLR* could be targeted for developing a treatment strategy for NAFLD with a minimum side effect to other human tissues.

The aim of the present study is two folds: 1) investigate the driving role of *PKLR* in NAFLD progression in multi-tissue context and 2) identify potential small molecular drugs for inhibition of *PKLR*. In this study, we investigated the critical role of the *PKLR* in NAFLD in a *Pklr* knock out (KO) mouse model and revealed the underlying molecular mechanisms associated with the preventive role of *Pklr* KO in multiple tissues. In addition, we repositioned a few small molecular drugs through a computational pipeline that may modulate expression level of *PKLR* and evaluated the top candidates in *in vitro* model. Finally, we evaluated two small molecular drugs with best performance in *in vitro* experiments in an *in vivo* rat study which follows a preclinical setup for promoting them into NAFLD clinical trials.

## Materials and Methods

Male C57BL/6J mice were used for mice experiment, and the *Pklr* KO mice as well as the wild-type ones are purchased from Applied StemCell company. All mice were housed at the University of Gothenburg animal facility (Lab of Exp Biomed) and supervised by university veterinarians and professional staff. Healthy Sprague Dawley male rats were obtained from Atatürk University Experimental Research Center (ATADEM, Erzurum Turkey) and used for the biosafety and efficacy were rat studies. The weights of the rats and comsumption of sucrose water of the rats are provided in Table S1.

WT HepG2 was used as *in vitro* model in this study and cultured by Roswell Park Memorial Institute (RPMI, R2405, Sigma Aldrich) 1640 Medium with 10% fetal bovine serum (FBS) and 1% penicillin – streptomycin (PS). To induce a steatosis model into WT HepG2 cells, high glucose Dulbecco’s Modified Eagle Media (DMEM, D0822/D5671, Sigma-Aldrich) containing 10% FBS and 1% PS was used as medium for one week, together with 10μg/mL insulin and 10μM T0901317.

The computational drug repositioning pipeline was adapted based on the concept introduced by Connectivity Map (Subramanian et al., 2017). The Pklr signature was determined by metaanalysis of three individual RNA-seq datasets, including two datasets from our previous study (Liu et al., 2019) and one for *in vivo* samples from GTEx.

Further experimental details are available in the Supplementary Methods. All raw RNA-sequencing data generated from this study can be accessed through accession number GSE. Codes used during the analysis are available on https://github.com/sysmedicine/pklr

## Results

### The effect of PKLR KO on liver

To systematically investigate the effect of pyruvate kinase liver isoform in multi tissue context, we fed four groups of 8-week-old *Pklr* KO and control C57Bl/6J mice (Ctrl) either with HSD or CHOW for eight weeks (Figure 1A).

**Figure 1.**
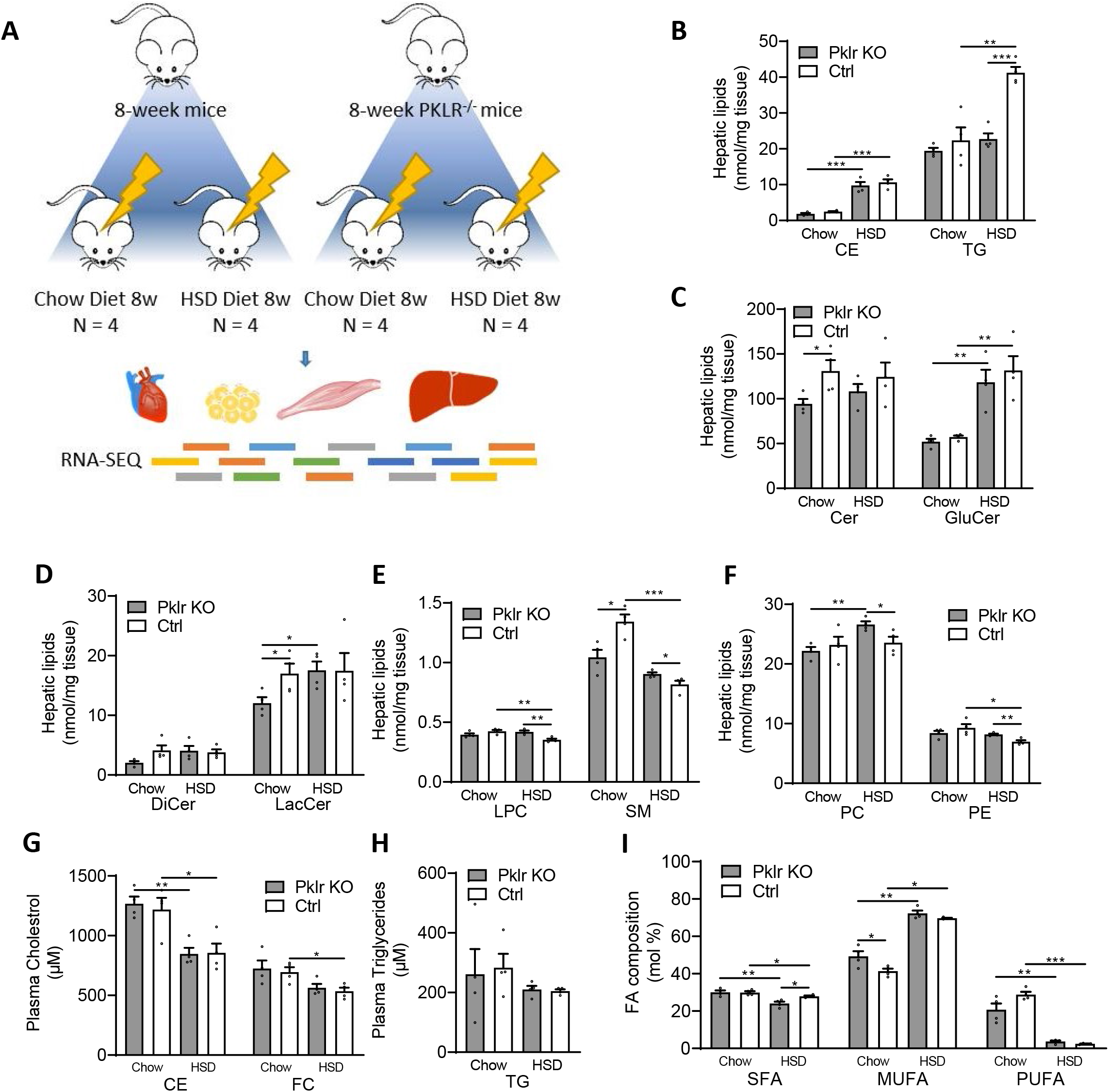
Analysis for liver lipidomics profile of 8-week wild-type and Pklr KO mice fed with CHOW and HSD. (A) Flow chart showing the experimental design. (B-F) Bar graphs showing the hepatic cholesterol ester (CE), triacylglyceride (TG), ceramide (Cer), glucosylceramide (GluCer), dihydroceramides (DiCer), lactosylceramide (LacCer), lysophosphatidylcholines (LPC), sphingomyelin (SM), phosphatidylcholines (PC) and phosphatidylethanolamine (PE) levels of mice in different conditions. (G-H) Bar graphs showing the plasma cholesterol ester (CE), free cholesterol (FC) and triacylglyceride (TG) levels of mice in different conditions. (I) Bar graph showing the composition of saturated fatty acids (SFA), mono unsaturated fatty acids (MUFA) and poly unsaturated fatty acids (PUFA) levels of mice in different conditions. *P<0.05, **P<0.01, ***P<0.001.

We carefully examined the liver and plasma lipid levels as well as the fatty acids composition in all four groups of mice (Figure 1B-I). We found that HSD-fed mice developed significant liver fat compared to CHOW-fed mice (Figure 1B). The *Pklr* KO mice were well tolerant of the HSD and *Pklr* KO HSD-fed mice had significantly lower liver TG levels than the Ctrl HSD-fed mice (Figure 1B). In addition, we found that the hepatic level of ceramide, lactosylceramide and sphingomyelin levels were significantly decreased in *Pklr* KO mice compared to the Ctrl Chow-fed mice (Figure 1C-E). We also found lysophosphatidylcholine, sphingomyelin, phosphatidylcholines and phosphatidylethanolamine are significantly increased in *Pklr* KO HSD-fed mice compared to the Ctrl HSD-fed mice (Figure 1E&F). No significant changes in plasma cholesterol or triglycerides we observed between Pklr KO and Ctrl mice (Figure 1G&H), and notably, a significant decrease of saturated fatty acids percentage in *Pklr* KO HSD-fed mice compared to the Ctrl HSD-fed mice (Figure 1I).

To systematically study how the *Pklr* KO prevent the development of liver fat in mice fed with HSD, we collected liver tissue samples from these four groups of mice and generated RNA-seq data. We firstly examined the hepatic *Fasn* expression and found that it is significantly elevated in Ctrl HSD-fed mice but not in the *Pklr* KO HSD-fed mice (Figure 2A). Our analysis indicated that *Pklr* KO inhibits the hepatic DNL in HSD-fed mice. We found that there is no significant difference in Fasn expression between the *Pklr* KO CHOW-fed mice and Ctrl CHOW-fed mice, and this is also supported by the measurement of comparable liver TG levels between these two groups of mice (Figure 1B).

**Figure 2.**
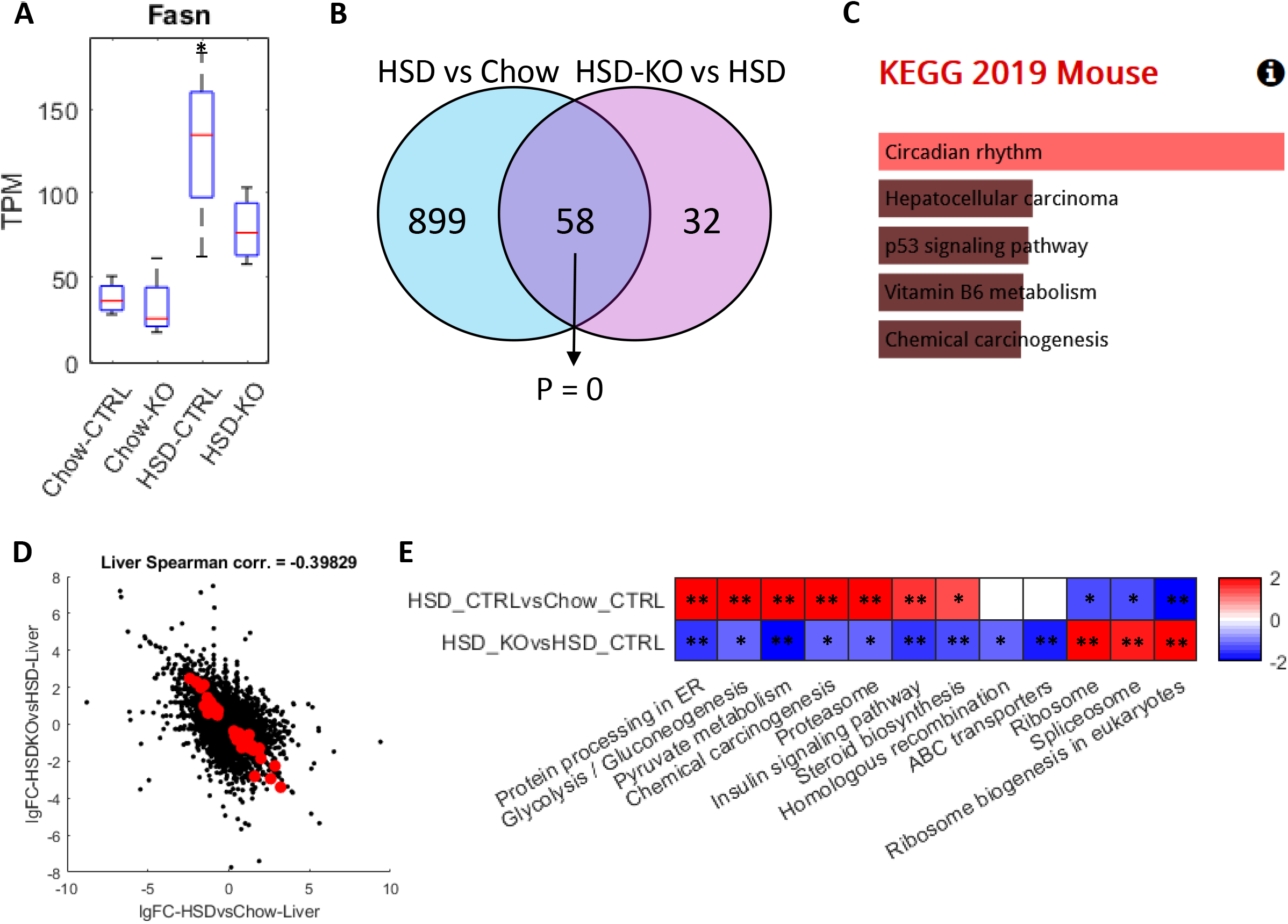
Analysis for liver transcriptomics profile of 8-week wild-type and Pkl KO mice fed with CHOW and HSD. (A) Box plot showing the expression of Fasn in different conditions, and the asterisks indicated the significant increase of Fasn expression compared to wild-type mice fed with CHOW (P < 0.05). (B) Venn-diagram showing the number and hypergeometric P value of overlapped DEGs between HSD Ctrl vs CHOW Ctrl and HSD Ctrl vs HSD KO in liver. (C) Barplot showing the significantly enriched KEGG pathways of the overlapped DEGs in Figure 2B. (D) Scatter plot showing the correlation between the log 2 fold changes of gene expression of all genes in HSD Ctrl vs CHOW Ctrl (x-axis) and HSD KO vs HSD Ctrl (y-axis), where the dots highlighted in red are the overlapped DEGs highlighted in Figure 2B. (E) Heatmap showing the differentially expressed pathways in the two comparisons, respectively, where each cell in the plot shows the minus log 10 P value if they are up-regulated, and positive log 10 P value if they are down-regulated (* P adj. <0.1; ** P adj. <0.05). Only pathways that are significantly differentially expressed in HSD KO vs HSD Ctrl (P adj. <0.1) are displayed and pathways without significant expression alterations (P adj. >0.1) are marked white.

Next, we focused on two comparisons, Ctrl HSD-fed vs Ctrl CHOW-fed and *Pklr* KO HSD-fed vs Ctrl HSD-fed mice and analyzed the transcriptomic changes between high liver fat vs normal as well as *Pklr* KO treatment vs high liver fat, respectively. We performed differential expression analysis and pathway enrichment analysis using KEGG pathways. We identified 957 DEGs between Ctrl HSD-fed vs Ctrl CHOW-fed mice and 90 genes in *Pklr* KO HSD-fed vs Ctrl HSD-fed mice (Table S2). Interestingly, we found that 58 DEGs are shared between these two groups and showed a significant transcriptomic change in the opposite direction (hypergeometric P = 0; Figure 2B). We performed functional enrichment analysis using these 58 overlapped DEGs and found that they are only significantly enriched in circadian rhythms (P adj. < 0.05; Figure 2C; Table S3), which suggests the *Pklr* KO reversed the dysregulation of circadian rhythms induced by HSD. Notably, the expression level of these DEGs in the circadian rhythms pathway is not significantly altered in the *Pklr* KO mice compared with the wild-type mice fed with CHOW (P adj. > 0.05; Table S2).

We analyzed the transcriptomic changes’ landscape and found an apparent negative correlation between the fold changes of the transcriptomic expression between these two comparisons (Spearman corr. = −0.398; Figure 2D). We also employed the PIANO toolbox (Varemo et al., 2013) to investigate the pathway level transcriptomic changes based on the KEGG pathways (Kanehisa et al., 2021). The pathway-level changes we identified in HSD vs CHOW included up-regulation of glycolysis, insulin signaling pathway, protein processing in ER, fatty acid metabolism, steroid and terpenoid metabolism and down-regulation of amino acid metabolism and transcription (Figure S1). Although less number of pathways showed significant alteration in the *Pklr* KO HSD-fed compared to Ctrl HSD-fed mice, the transcriptomic expression of several these pathways, such as protein processing in ER, glycolysis, insulin signaling pathway, steroid biosynthesis, ribosome and spliceosome pathways, are significantly reversed after *Pklr* KO (Figure 2E).

### Investigating NAFLD progression and Pkl KO effect in other metabolic tissues

It has been reported that the progression of NAFLD is associated with extrahepatic tissues, such as WAT, muscle and heart tissues (Haas et al., 2016). Hence, it is vital to analyse the global transcriptomics changes in multi-tissue context to study the molecular differences during NAFLD progression and *Pklr* KO. We obtained WAT, muscle and heart tissue samples from the 8-week Ctrl and *Pklr* KO mice fed with both CHOW and HSD and generated RNA-Seq data. First, we performed differential expression analysis between Ctrl HSD-fed vs Ctrl CHOW-fed and *Pklr* KO HSD-fed vs Ctrl HSD-fed mice in these three tissues and compared them with the DEGs in the liver (Figure 3A; Table S2).

**Figure 3.**
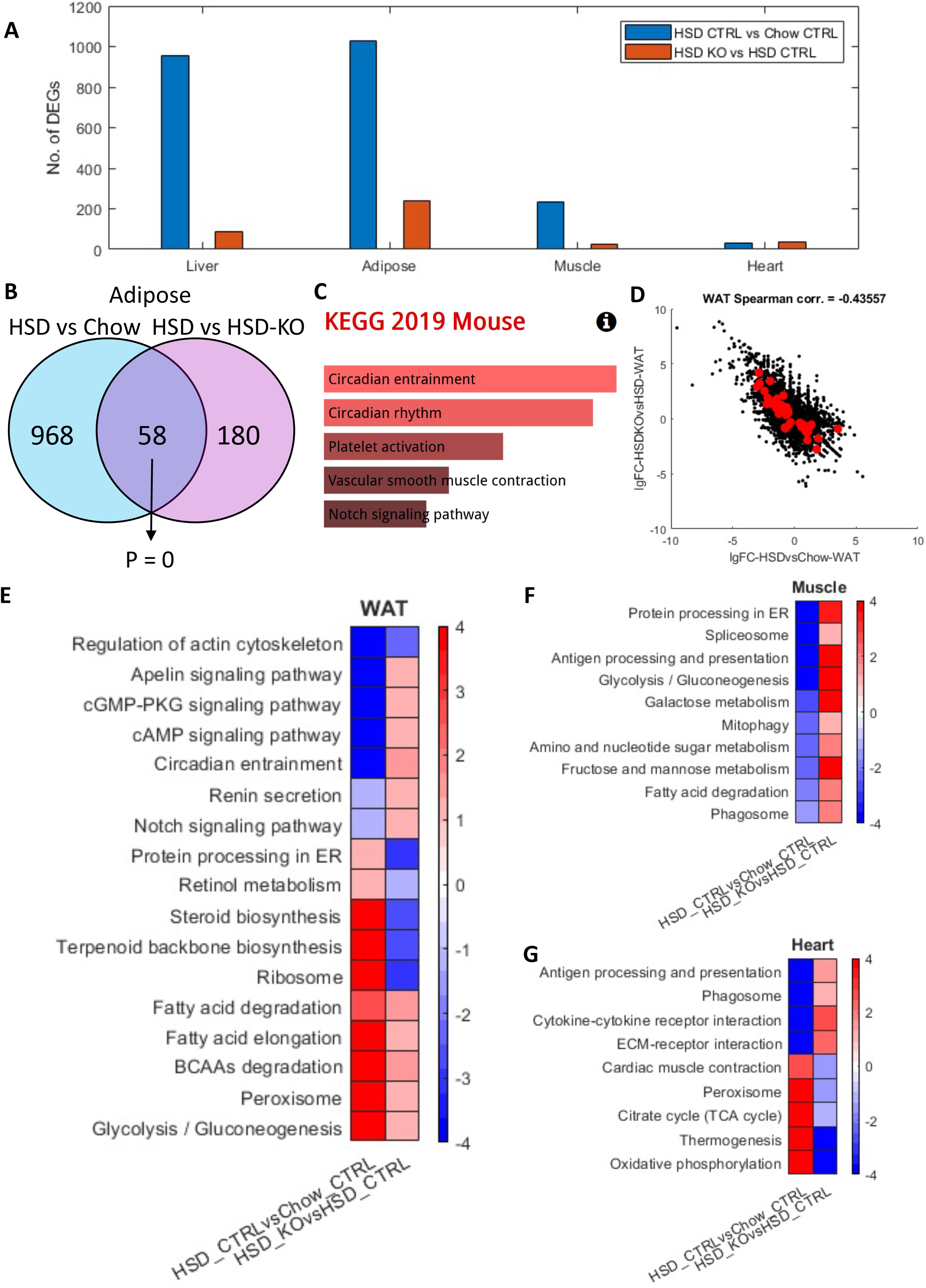
Analysis for transcriptomics profile of muscle, WAT and heart obtained from 8-week wild-type and Pkl KO mice fed with different CHOW and HSD. (A) Bar graph showing the number of DEGs in indicated comparisons liver, WAT, muscle and heart. (B) Venn-diagram showing the number and hypergeometric P value of overlapped DEGs between HSD Ctrl vs CHOW Ctrl and HSD Ctrl vs HSD KO in WAT. (C) Barplot showing the significantly enriched KEGG pathways of the overlapped DEGs in Figure 4B. (D) Scatter plot showing the correlation between the log 2 fold changes of gene expression of all genes in HSD Ctrl vs CHOW Ctrl (x-axis) and HSD KO vs HSD Ctrl (y-axis), where the dots highlighted in red are the overlapped DEGs highlighted in Figure 4B. (E-G) Heatmaps showing the differentially expressed pathways in the two comparisons in WAT, muscle and heart, respectively, where each cell in the plot shows the minus log 10 P value if they are up-regulated, and positive log 10 P value if they are down-regulated (* P adj. <0.1; ** P adj. <0.05). Only pathways that are significantly differentially expressed in both HSD KO vs HSD Ctrl and HSD ctrl vs CHOW Ctrl (P adj. <0.1) are displayed.

Interestingly, we observed the largest number of DEGs in WAT among all comparisons (1026 and 238, respectively), followed by the muscle (236 and 5, respectively) and heart (32 and 2, respectively). We checked the overlapped DEGs between the two comparisons (Ctrl HSD-fed vs Ctrl CHOW-fed and *Pklr* KO HSD-fed vs Ctrl HSD-fed) in WAT which showed the greatest transcriptomic changes in terms of DEGs number and found that 58 genes are significantly reversely altered in these two comparisons (hypergeometric P = 0; Figure 3B), which is very similar to what we observed in the liver. In addition, although not statistically significant, the top enriched KEGG pathways of these 58 genes are circadian entrainment and circadian rhythm (FDR = 0.17; Table S4), which is also in excellent agreement with the 58 reversed changed DEGs identified in the liver (Figure S2). This might indicate that circadian regulation played an important role in the whole-body response to *Pklr* KO. Moreover, we observed a negative correlation between the fold changes of the gene expressions between these two comparisons (Spearman corr. = −0.436; Figure 3D). We also investigated the changes in muscle and heart tissues. Notably, the DEGs are also significantly reversely overlapped in muscle and heart tissues (hypergeometric P < 0.05) and their gene expression changes showed negative correlations as well (Figure S3). Taken together, we observed that the hepatic *Pklr* KO partly reversed the transcriptomic changes in extrahepatic tissues induced by HSD.

Next, we investigated the pathway level changes using the DEGs in all three tissues using PIANO and KEGG pathways in the same way as in the liver tissue. In WAT, we observed significant up-regulation of protein processing in ER, ribosome, retinol metabolism and steroid and terpenoid backbone biosynthesis pathways and significant down-regulation of several signaling pathways, such cAMP signaling, notch signaling, cGMP-PKG signaling, apelin signaling pathways, between Ctrl HSD-fed vs Ctrl CHOW-fed mice. These pathways are significantly changed in the opposite direction between *Pklr* KO HSD-fed vs Ctrl HSD-fed mice (Figure 3E). These pathways with the reversed gene expression changes indicated the changes in WAT in response to *Pklr* KO. For instance, the reverse of steroid and terpenoid backbone biosynthesis pathways implicated the potential link between *Pklr* and whole-body steroid hemostasis. Notably, in both Ctrl HSD-fed vs Ctrl CHOW-fed and *Pklr* KO HSD-fed vs Ctrl HSD-fed mice, glycolysis, peroxisome, BCAAs degradation, fatty acid degradation and elongation pathways are significantly up-regulated in WAT, while regulation of actin cytoskeleton pathway is significantly down-regulated. The gene expression changes in these pathways are enhanced rather than reversed with *Pklr* KO, suggesting the complementary response of these pathways to the hepatic gene expression changes. This is exemplified by the enhanced up-regulated glycolysis pathway, as its up-regulation in Ctrl HSD-fed vs Ctrl CHOW-fed mice is probably caused by the excessive amount of glucose intake in the diet. The up-regulation in *Pklr* KO HSD-fed vs Ctrl HSD-fed mice implicated that the decreased glucose intake in the liver in response to *Pklr* KO is complemented by the further enhanced glycolysis in WAT.

In the muscle tissue, we observed that the *Pklr* KO reversed the down-regulation of metabolic pathways, including glycolysis, galactose metabolism, fructose and mannose metabolism, amino and nucleotide sugar metabolism and fatty acid degradation, and other pathways such as protein processing in ER, antigen processing and presentation, mitophagy, phagosome and spliceosome pathways. In the heart tissue, the *Pklr* KO reversed the down-regulation of antigen processing and presentation, phagosome, cytokine-cytokine receptor and ECM-receptor interaction pathways and attenuated the expression of the up-regulated pathways, including cardiac muscle contraction, peroxisome, TCA cycle, thermogenesis, oxidative phosphorylation. Notably, no pathway in muscle or heart showed enhanced transcriptomic changes after *Pklr* KO in response to HSD. In summary, we concluded that the *Pklr* KO reversed the transcriptomic changes induced by HSD in extrahepatic tissues, and the affected pathways are mostly related to protein processing, carbohydrate metabolism, fatty acids metabolism, and oxidative phosphorylation as well as some immune and cell recognition associated pathways.

### Drug repositioning for inhibition of Pklr in vitro

Based on the *in vivo* experiment in this study, we concluded that *PKLR* is a promising drug target to treat NAFLD. Therefore, we performed drug repositioning to identify a therapeutic agent that can target *PKLR* and could be tested in clinical trials. In this context, we developed a computational pipeline for repositioning of small molecule drugs that can effectively inhibit *PKLR* using the data in the Connectivity Map (Subramanian et al., 2017) (Figure 4A). In brief, we analysed human liver tissue RNA-Seq data obtained from 110 subjects in the GTEx database and HepG2 cell line RNA-Seq data where we inhibited/over-expressed *PKLR*, and identified a *PKLR*-related consensus gene signature (Table S5). We compared this consensus gene signature with the gene expression profiles of more than 1800 compound-perturbed-HepG2 from the LINCS L1000 database and identified the potential inhibitors that show similar gene expression profile (see method). As a result, we selected the 10 drugs with the most negative expression correlation with the *PKLR*-related consensus gene signature and identified these drugs as potential *PKLR* inhibitors (Table S6).

**Figure 4.**
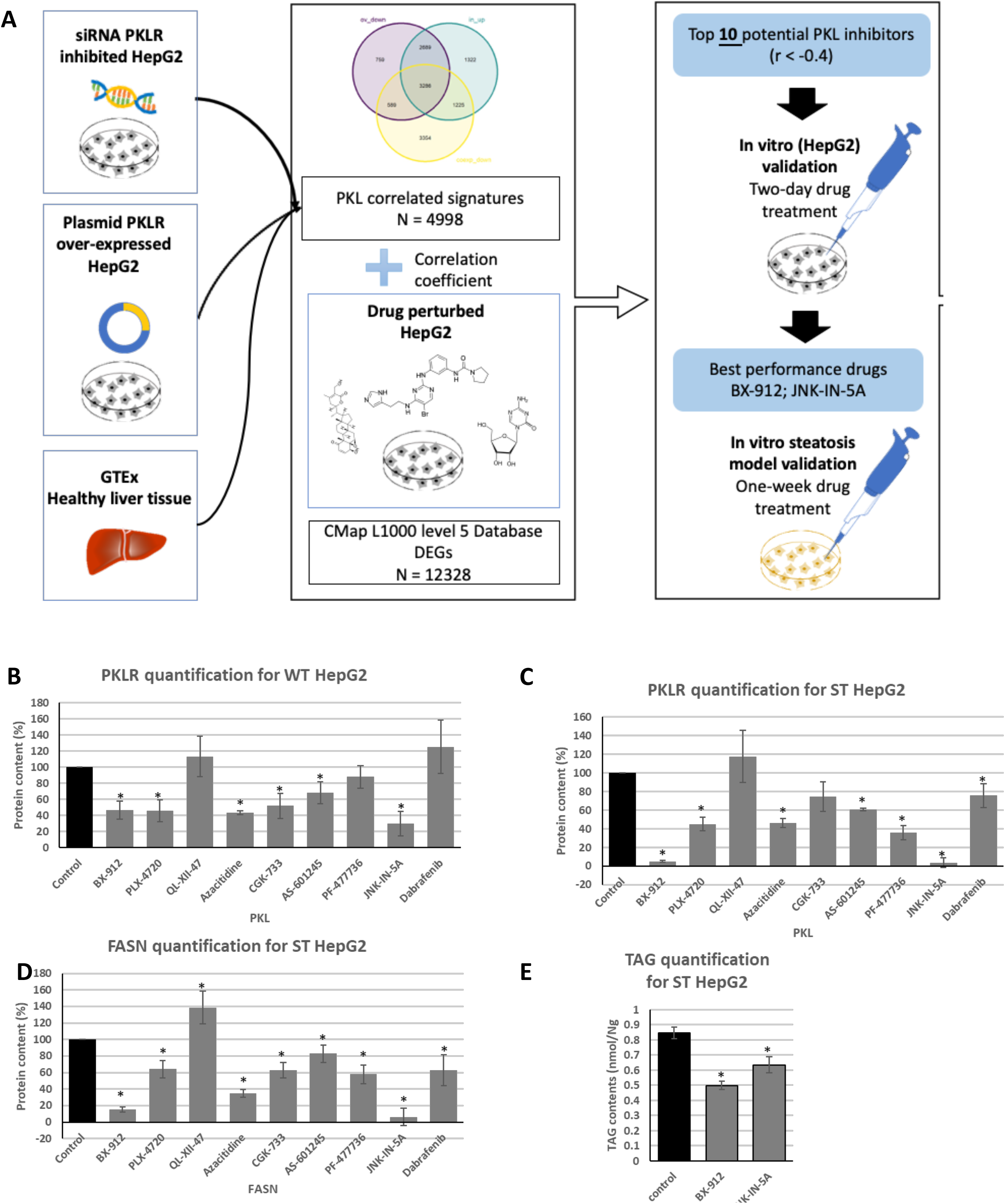
Drug repositioning application and in vitro validation of potential Pklr inhibitors. (A) Workflow. (B-D) Bar plot showing the drug effects on protein expression of HepG2 and steatosis-induced HepG2, normalized by ß-actin. (E) Bar plot showing the drug effects on triglyceride contents of steatosis-induced HepG2. The results were normalized by cell viability. Results were normalized by protein content. (G) Bar plot showing the results of western blot of liver biopsy, normalized by ß-actin. (*: p-value < 0.05; t-test)

We evaluated all 10 drugs by performing a literature review (Table S7) and found some of these drugs have been associated with fatty acid metabolism. For instance, Azacitidine, which was reported as a DNA methyltransferase inhibitor, has been approved by the US FDA for the treatment of all subtypes of myelodysplastic syndromes (MDS) (Muller and Florek, 2010). It has also been found that the drug reduces the expression level of a list of genes controlling fatty acid biosynthesis *in vitro* and *Pcsk9* (Poirier et al., 2014). In the high-glucose-cultured HepG2 cell model, the agent decreases the expression level of another key DNL gene, *Fasn* (Hosseini et al., 2020). Among the other 9 compounds, six of them turned out to be kinase inhibitors, indicating their potential effect on *PKLR*. Additionally, AS-601245 and JNK-IN-5A were reported as the kinase inhibitor of c-Jun N-terminal kinases (JNK) (Cerbone et al., 2012).

To determine the inhibitory effects of drugs on the expression level of *PKLR*, HepG2 was selected as the *in vitro* model for coherence. We purchased nine drugs (BX-912, PLX-4720, QL-XII-47, Azacitidine, CGK-733, AS-601245, PF-477736, JNK-IN-5A, Dabrafenib) from the list that are commercially available and treated HepG2 cells with these drugs for two days with different dosages due to their potential cellular toxicities (Table S8), and then measured the *PKLR* protein level by western blotting. The protein expression levels of *PKLR* were compared between the drug-treated and Ctrl group after normalizing with β-actin. As shown in Figure 4B, six of the nine tested drugs significantly down-regulated the protein expression of *PKLR* (P < 0.05).

To further analyse the effect of these drugs on lipid accumulation via the DNL pathway, we induced steatosis to HepG2 by treating the cells with high glucose medium (DMEM), insulin and T0901317 (see Method) (Hansmannel et al., 2006). After 7 days of drug treatment, eight of the nine tested drugs significantly decreased the protein expression level of PKLR (P < 0.05; Figure 4C). Six of the nine tested drugs also significantly reduced the protein expression level of *FASN* (P < 0.05; Figure 4D). Specifically, BX-912 and JNK-IN-5A treated groups showed a dramatic reduction in the protein expression level of both *PKLR* and *FASN* in the steatosis model. Therefore, we investigated the effects of these two drugs on the *de novo* synthesis of lipids based on the steatosis-induced HepG2 model. After 7 days of drug treatment, both drugs significantly decreased the content of TGs in the HepG2 cells (Figure 4E). Taken together, we showed that BX-912 and JNK-IN-5A effectively alleviate the protein expression of *PKLR* and *FASN* as well as TG levels in HepG2 cell lines, and thus, we decided to perform *in vivo* evaluations to test the effect of these two drugs.

### The treatment of rats with the therapeutic agents

To evaluate the potential toxicity of BX-912 and JNK-IN-5A in animals, we raised three groups of 12-week-old rats (n = 5), fed the first group with CHOW (Ctrl), the second group with CHOW plus BX-912 (30 mg/kg) and the third group with CHOW plus JNK-IN-5A (30 mg/kg) for seven days (Figure 5A). After 7 days, all rats were sacrificed, and tissue samples from major organs and blood samples were collected for biosafety evaluation. We analysed the frequencies of micro nucleated polychromatic erythrocytes (MNPCEs) in the blood samples, and found that the frequencies of MNPCEs in drug-treated groups were not statistically different (p>0.05) when compared to Ctrl animals. Our analysis indicated that both tested agents exhibited non-genotoxic potential (Figure 5B; Table S9). In addition, we performed histopathological examinations using the tissue samples obtains from the rats and found that kidney, small intestine, large intestine, stomach, heart, pancreas, muscle tissues and liver from all three rat groups exhibited normal histological structures (Figure 5C and D). Moreover, we performed immunohistochemical (IHC) examination on the liver tissues from the rats and observed negative 8-OH-dG expression in all three groups. Therefore, we concluded that both BX-912 and JNK-IN-5A have no side effects on the rats.

**Figure 5.**
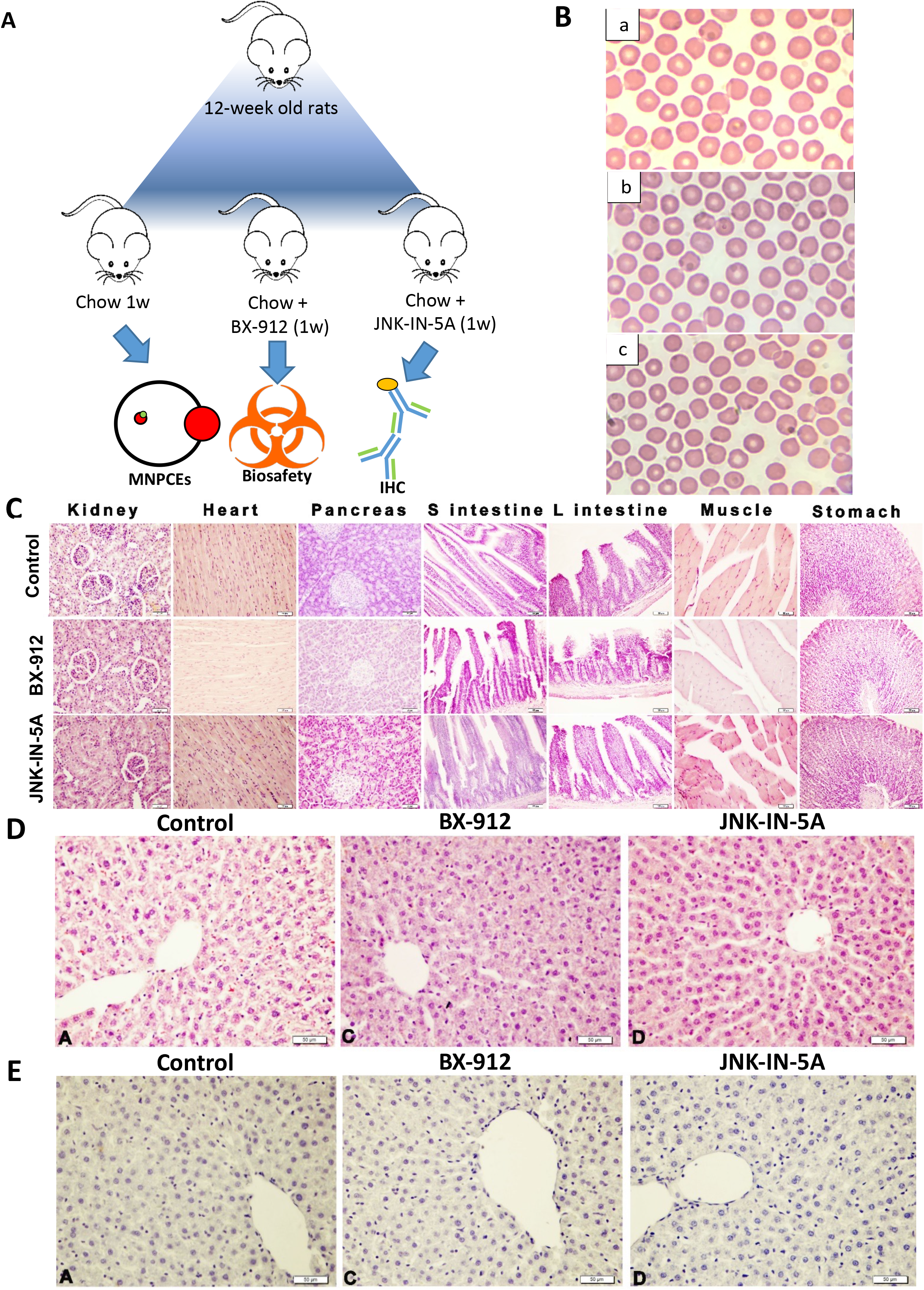
In vivo biosafety examination of the potential Pklr inhibitors. (A) Workflow. (B) Light micrograph of rat blood smear obtained from (a) control group (b) treated with 30 mg/kg BX-912 (c) treated with 30 mg/kg JNK-512 for 7 days (Giemsa stain, 1,000x magnification). (C) Histological images from major organs of the indicate rat groups where Control, BX and JNK represents rats fed with chow diet, chow diet plus BX-912 and chow diet plus JNK-IN-5A, respectively. (D) Histological images from liver tissue samples from respective rat groups where arrows and arrowheads respectively indicate degeneration and steatosis. Bar: 50μm. (E) IHC images with staining showing 8-OH-dG expression indicated by arrowheads in liver tissues from corresponding rat groups.

Next, we investigated the efficacy of these two drugs in a steatosis rat study by feeding the animals with HSD. We raised five groups of 12-week-old rats (n = 5), fed the first group with CHOW diet and the remaining four groups with HSD for two weeks (Figure 6A). At the end of week 2, we sacrificed the CHOW-fed rats (n = 5) and HSD-fed rats (n = 5) for histopathological examination and confirmed the HSD fed rats already developed hepatic steatosis.

**Figure 6.**
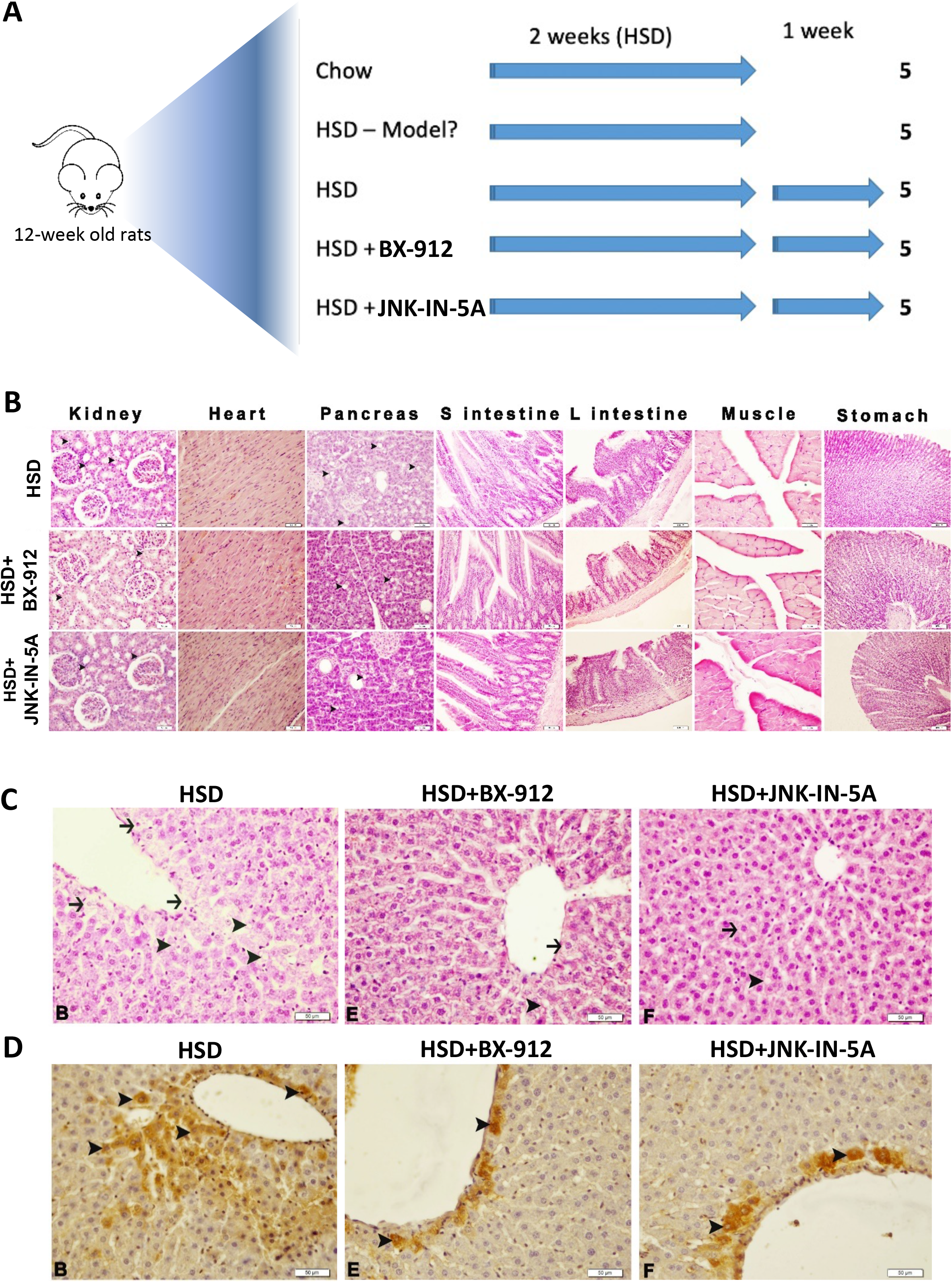
In vivo efficacy experiment for the potential Pklr inhibitors. (A) Study design of the experiment. (B) Histological images from liver tissue samples from respective rat groups where arrows and arrowheads respectively indicated degeneration and steatosis. Bar: 50μm. (C) IHC images with staining showing 8-OH-dG expression indicated by arrowheads in liver tissues from corresponding rat groups. (D) Bar plot illustrating the triglyceride contents of liver biopsy of drug-treated mice. (E) Bar plot showing the hepatic protein expression of indicated proteins quantified by western blot, normalized by ß-actin. (*: p-value < 0.05; t-test)

After securing the development of the liver steatosis model, we fed the third group with HSD (HSD 3w), the fourth group with HSD plus BX-912 (30 mg/kg) and the fifth group with HSD plus JNK-IN-5A (30 mg/kg) for 7 days. After 7 days, all three group of rats were sacrificed, and tissue samples from the liver and other major organs and blood samples were collected for efficacy examination. We performed a histopathological and IHC analysis on the liver samples and other major organs obtained from the three rat groups. We observed a standard histological structure on the small intestine, large intestine, stomach, heart and muscle tissues in the examination for drug-treated groups and found no toxic effect of the small molecule drugs on these tissues (Figure 6B). In the rats from HSD 3w group, severe degeneration was detected in the renal tubule epithelium and severe fattening was detected in the parenchyma cells in the pancreatic tissues. Interestingly, we detected only mild degeneration in renal tubule epithelium and mild fattening in the parenchyma cells of pancreas tissues in drug-treated groups. In addition, we observed severe degeneration and steatosis in hepatocytes in liver tissues of rats from the HSD 3w group, mild degeneration and moderate steatosis in hepatocytes in liver tissues of the HSD-fed rats treated with BX-912, while observed only mild degeneration and steatosis in the liver tissues of the HSD-fed rats treated with JNK-IN-5A (Figure 6C; Table S10). Moreover, strong 8-OH-dG expression was observed in hepatocytes in liver tissues of the rats from HSD 3w group, and the 8-OH-dG expression is significantly decreased in drug-treated groups compared to the HSD 3w rats (Figure 6D; Table S10).

## Discussions

In our mice experiment, we observed that *Pklr* KO in liver could prevent the HSD-induced hepatic steatosis and functional alterations at transcriptomic level. We observed that the mRNA expression level of glycolysis, steroid biosynthesis and insulin signaling pathways are enhanced in livers of HSD-fed mice and attenuated *Pklr* KO HSD-fed mice. Enhancement of glycolysis and insulin resistance is a hallmark of NAFLD (Fabbrini et al., 2010), and the inhibition of these two pathways in *Pklr* KO mice highlighted the potential role of *Pklr* as a therapeutic target for NAFLD treatment. In fact, a recent study transfected shRNA and plasmid to respectively inhibit and over-express *Pklr* in mice liver, and demonstrated that *Pklr* is causal in developing insulin resistance (Chella Krishnan et al., 2021). In addition, the reverse effect of the steroid biosynthesis suggested a potential link between hepatic *Pklr* and steroid hormone homeostasis, known to be involved in the development of NAFLD (Charni-Natan et al., 2019).

Our study also for the first time reported the *Pklr* KO effect in extrahepatic tissues, and our results provided valuable insights about the biological function of *Pklr* in the whole-body context. We found that the most responsive extrahepatic tissue is WAT. The pathways showed significant transcriptomic alteration are very similar to those observed in liver, including glycolysis, steroid biosynthesis, protein processing in ER and ribosome pathways, implicated a synergetic regulation between liver and WAT. Interestingly, central circadian regulators also showed similar alterations based on transcriptomics data, which may implicate that *Pklr* induces the transcriptomic changes in WAT via synchronization of circadian rhythms. It is known that circadian clock includes glucose sensor and participates in the orchestration of glucose homeostasis (Asher and Schibler, 2011), and it has been reported that interruption of circadian rhythms might be an important contributor to the development of insulin resistance (Stenvers et al., 2019). In addition, a previous study reported the liver specific knockout of *Bmal1*, which is the key circadian regulator, disrupted the diurnal rhythms of hepatic *Pklr* expression (Udoh et al., 2018). Previous studies and our analysis may indicate the regulatory effect between the key circadian regulator and *Pklr* in the liver. In this context, it is possible that the *Pklr* KO could directly affect the circadian rhythm in liver. Moreover, both liver and WAT have autonomous circadian clocks which are regulated partly by glucose intake, and it is very likely for liver to synchronize its circadian clock with the WAT one via controlling glucose and lipid metabolism (Stenvers et al., 2019). Taken together, we speculate that the excessive intake of glucose in HSD-fed mice disrupts the circadian regulation and induces insulin resistance in liver and WAT. The knockout of *Pklr* inhibits the glycolysis in liver and restores the circadian rhythms in the liver and WAT.

To identify a small molecule drug that could potentially be used to treat NAFLD via inhibition of *Pklr*, we established a computational drug repositioning pipeline and found 10 candidate drugs that could potentially targeting *Pklr*. Unlike traditional method, the identified drugs were meant to decrease the expression of the target gene rather than its activity. We validated the inhibitory effect on *Pklr* expression as well as lipid accumulation of the six predicted drugs in an *in vitro* steatosis model. In addition, the two most effective small molecules drugs, namely BX-912 and JNK-IN-5A, were tested in rat animal study with a preclinical setup. Both drugs passed biosafety experiment in rat without causing noticeable side effect in the animals. In addition, we observed that both of these drugs significantly decreased liver fat and attenuated oxidative stress and degeneration of renal tubule epithelium and fattening in the parenchyma cells of pancreatic tissues in the rats, implicating a wholebody improvement of the rats which agrees well with what we observed in the multi-tissue transcriptomic analysis. These results highlighted the efficiency and power of computational drug repositioning pipeline.

A potential limitation in this study is that we used rat as *in vivo* model for validation of the drug repositioning instead of mouse which was used earlier in this study. Ideally, the same animal model should be used throughout the study, but we selected rat model for evaluation of the drug effect as it is a more commonly used preclinical model for biosafety and drug efficacy tests. Considering that the increase hepatosteatosis and inflammation (oxidative stress) were observed also in the HSD-fed rats, we concluded that the HSD fed rats could serve as a proper model for studying NAFLD as well as evaluating the drug effects.

In summary, our study demonstrated that knockout of *Pklr* prevent HSD-fed mice from developing liver fat by effecting liver, WAT, muscle, and heart tissues. We showed that the *Pklr* KO could reverse the HSD induced pathway transcriptomic changes and observed that these changes could be regulated via metabolically synchronized circadian rhythms. We also identified two small molecule drugs, which inhibit the *Pklr* expression and attenuate liver fat *in vivo*. Therefore, our study provided biological insights about the effect of *Pkl* KO *in vivo*, and presented the development of a promising therapeutic solution for treatment of NAFLD patients, which should be evaluated in future clinical trials.

## Acknowledgement

This work was financially supported by ScandiEdge Therapeutics and Knut and Alice Wallenberg Foundation.

The computations and data handling were enabled by resources provided by the Swedish National Infrastructure for Computing (SNIC) at UPPMAX, partially funded by the Swedish Research Council through grant agreement no. 2018-05973.

## SUPPORTING INFORMATION

Supporting Information includes Supplemental Appendix, 3 Supplementary Figures and 10 Supplementary Tables.

## Supplementary Tables

**Table S1.** Body weights of rats in biosafety and efficiency experiments as well as sucrose consumption in the efficiency experiment.

**Table S2.** Result of differential expression analysis for all relevant comparisons of liver, white adipose tissue, muscle and heart in the 8-week mouse experiment.

**Table S3.** Enriched KEGG pathways of the overlapped DEGs between HSD KO vs. HSD Ctrl and HSD Ctrl vs. Chow Ctrl in liver in the 8-week mouse experiment.

**Table S4.** Enriched KEGG pathways of the overlapped DEGs between HSD KO vs. HSD Ctrl and HSD Ctrl vs. Chow Ctrl in white adipose tissue in the 8-week mouse experiment.

**Table S5.** The consensus *PKLR* signature genes and *PKLR*-correlated signature genes specifically obtained from in vitro siRNA/plasmid experiment and GTEx human liver dataset as well as the genes overlapped with CMap genes.

**Table S6.** Correlation between all drug signatures in HepG2 and the Pklr consensus signature and the top 10 negatively correlated drugs.

**Table S7.** Detailed information and literature review for the top 10 selected drugs.

**Table S8.** Dosage for all drug treatment in the in vitro experiment.

**Table S9.** Frequency of MN in experimental groups

**Table S10.** Scoring of histopathological and immunohistochemical findings in liver tissues

## Supplementary Figures

**Figure S1.**
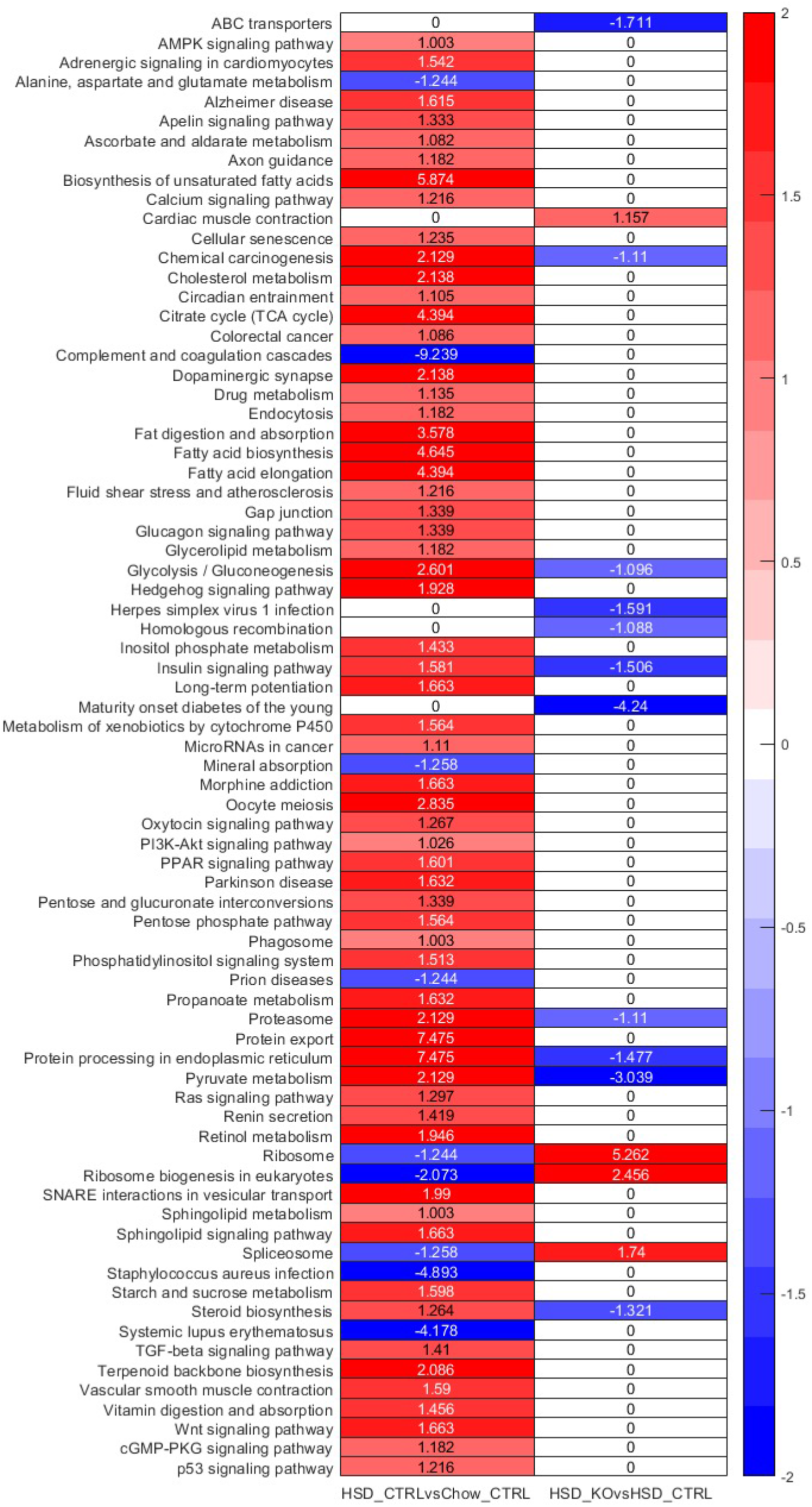
Heatmap showing the differentially expressed pathways in HSD Ctrl vs CHOW Ctrl and HSD KO vs HSD Ctrl, respectively, where each cell in the plot shows the minus log 10 P value if they are up-regulated, and positive log 10 P value if they are down-regulated. Only pathways that are significantly differentially expressed in either comparison (P adj. <0.1) are displayed and pathways without significant expression alterations (P adj. >0.1) are marked white.

**Figure S2.**
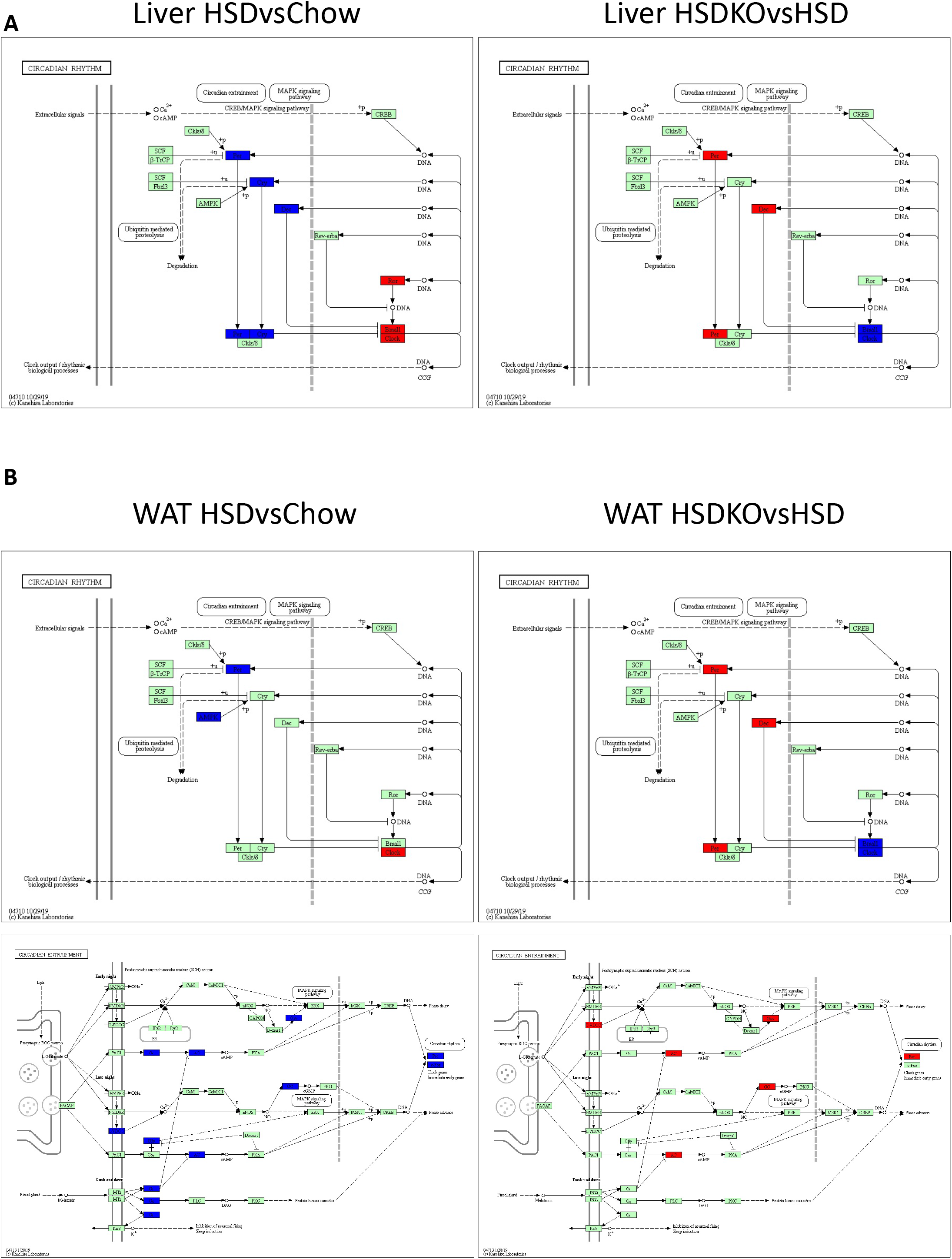
(A) Pathway figure generated by KEGG mapper highlighting the up- (RED) and down-regulated (blue) genes in circadian rhythm pathway in livers in HSD Ctrl vs CHOW Ctrl and HSD KO vs HSD Ctrl. (B) Pathway figure generated by KEGG mapper highlighting the up- (RED) and down-regulated (blue) genes in circadian rhythm and circadian entrainment pathways in white adipose tissue in HSD Ctrl vs CHOW Ctrl and HSD KO vs HSD Ctrl.

**Figure S3.**
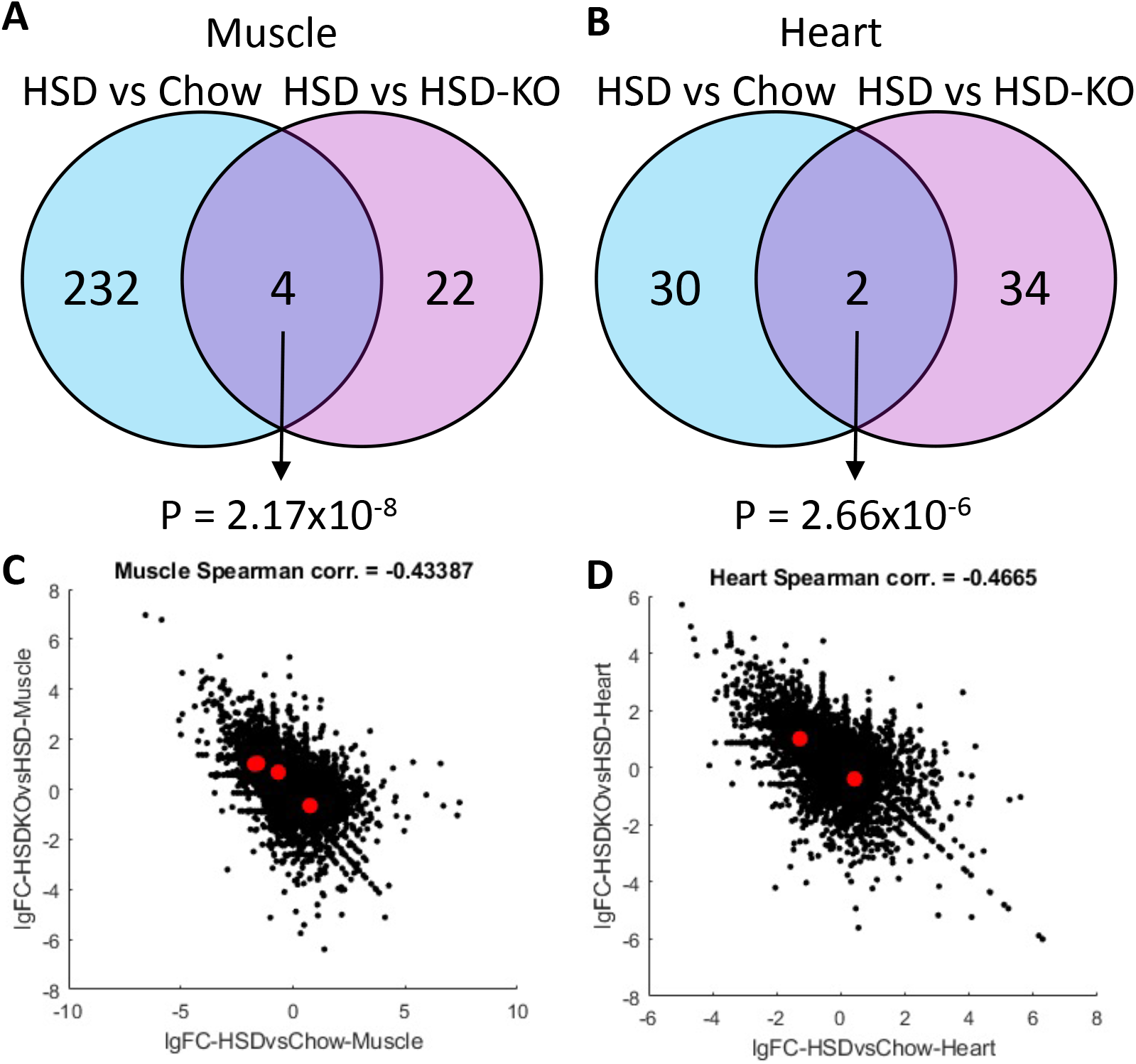
(A-B) Venn-diagram showing the number and hypergeometric P value of overlapped DEGs between HSD Ctrl vs CHOW Ctrl and HSD Ctrl vs HSD KO in muscle and heart. (C-D) Scatter plot showing the correlation between the log 2 fold changes of gene expression of all genes in muscle and heart in HSD Ctrl vs CHOW Ctrl (x-axis) and HSD KO vs HSD Ctrl (y-axis), where the dots highlighted in red are the overlapped DEGs highlighted in Figure 3D.

## Mouse animal studies

Eight male C57BL/6J mice wild-type mice and eight Pklr^-/-^ mutant mice were fed a standard mouse chow diet (Purina 7012, Harlan Teklad) and housed in a 12-h light–dark cycle. The *Pklr* KO mice as well as the wild-type ones are purchased from Applied StemCell company. The company generated a knockout (KO) model in mouse *Pklr* gene. The model specifies a Guanine (G) deletion immediate downstream of the ATG start codon in mouse *Pklr* L^-^ isozyme, not R-isozyme. (Supplementary report 1 & 2). From the age of 8 weeks, both the wild-type and mutant mice were then divided into two groups of 4 mice fed with chow diet, high-sucrose diet for another 8 weeks, respectively. At the age of 16 weeks, all mice are sacrificed and necropsies from liver, muscle, WAT and heart tissue were taken for RNA sequencing and lipid quantification.

All mice were housed at the University of Gothenburg animal facility (Lab of Exp Biomed) and supervised by university veterinarians and professional staff. The health status of our mice is constantly monitored according to the rules established by the Federation of European Laboratory Animal Science Associations. The experiments were approved by the Gothenburg Ethical Committee on Animal Experiments.

## Rat animal studies

### In vivo biosafety studies

Healthy Sprague Dawley male rats (12 weeks old, 260-300 g) were obtained from Atatürk University Experimental Research Center (ATADEM, Erzurum Turkey) and housed under standard environmental conditions (temperature 20 °C–25 °C, humidity 50 ± 20% and 12-hour light/dark diurnal cycle). Food and water were available ad libitum. Rats were randomly separated into one control group and two experimental groups (30 mg/kg/day BX-912 or JNK-IN-5A, n=5). BX-912 and JNK-IN-5A were dissolved in DMSO and were orally administrated using oral gavage for 7 days. The control group were received the same amount of vehicle solution only. The weights of the rats are provided in Table S1.

After 7 days, rats were anaesthetized and blood samples were collected from the abdominal aorta for biochemical and hematological analysis. The plasma was separated and stored at −80 °C for further analysis. Major organs such as: heart, kidney, liver, muscle, adipose, intestine (duodenum, ileum, jejunum), pancreas, colon, and stomach were collected from all the animals for further analysis. Selected organs were fixed in 10% neutral buffered formalin for histopathological examination.

In addition to sampling for biochemical and hematological analysis, one drop of blood samples was obtained from the animals for evaluating micronucleus frequencies as genotoxicity endpoint at the end of the experiments. Three different smears of each rat were prepared using pre-coded and cold slides, then the slides air-dried at room temperature, and fixed in ethanol for 10 min, and stained with Giemsa. The frequencies of micro nucleated polychromatic erythrocytes (MNPCEs) were analyzed according to previously suggested approach using a light microscope. The incidence of MN-PCEs was determined by scoring a total of 3000 PCE per animal and results were expressed as % MN (Solórzano-Meléndez et al., 2021).

### In vivo efficacy studies

After 1-week acclimation period, rats were randomly separated into two groups: control group (n=5) and high sucrose group (n=20). The control group was fed with a standard diet while the others consumed high sucrose diet (HSD). Sucrose was purchased from Sigma Chemical Co. (St Louis, MO, USA) and supplied at the dose of 10% in the drink water for two weeks. The consumption of sucrose water and weights of the rats are provided in Table S1.

After 2 weeks, five rats in the HSD group were randomly sacrificed for conformation of NAFLD model. Then, the remaining rats in the HSD group were separated into 3 sub-groups (n=5): HSD group, HSD plus BX-912 group and HSD plus JNK-IN-5A group. BX-912 and JNK-IN-5A were dissolved in DMSO and were orally administrated using oral gavage for one week (30 mg/kg/day). HSD group were treated with vehicle solution only.

At the end of the experiment, rats were sacrificed after being anesthetized. Blood samples were collected from the abdominal aorta and centrifuged at 8000 rpm for 15 min at 4 °C. The plasma was separated and stored at −80 °C until further analysis. Other internal organs, including the heart, adipose tissues, liver, kidney, muscle, intestine (duodenum, ileum, jejunum), pancreas, colon and stomach were immediately removed and then snap frozen in liquid nitrogen and stored at −80°C; the liver was also fixed in 10% formaldehyde for histopathological examination.

The biosafety and efficacy studies were approved by The Ethics Committee of Atatürk University, and all experiments were carried out in accordance with relevant guidelines and regulations for the care and use of laboratory animals. Animal Experiments Local Ethics Committee of Atatürk University approved the experimental procedure described in this study (Approval Date: 27.08.2020; Approval Number: 42190979-000-E.2000208344).

### Histolopathological examination

The liver tissues were fixed in 10% buffered formaldehyde solution. After fixation, the tissues were passed through graded alcohol and xylene series and embedded in paraffin blocks. 5 micrometer thick sections were taken serially from the paraffin blocks. Histopathological changes were evaluated by performing hematoxylin eosin staining on the sections taken. Sections were evaluated according to histopathological findings as none (-), mild (+), moderate (++) and severe (+++).

### Immunohistochemical examination

After deparaffinization, the slides were immersed in antigen retrieval solution (pH6.0) and heated in microwave for 15 min to unmask the antigens. The sections were then dipped in 3% H2O2 for 10 min to block endogenous peroxidase activity. Protein block was dropped onto the sections, washed with PBS for 10 min. Primary antibodies (8-OH-dG, Cat No: sc-66036, Diluent Ratio: 1/100, Santa Cruz) were prepared and applied according to the usage conditions. Expose mouse and rabbit specific HRP/DAB detection IHC kit were used as follows: sections were incubated with goat anti-mouse antibody, then with streptavidin peroxidase, and finally with 3,3’ diamino benzidine + chromogen. Slides were counter stained with hematoxylin. Immunoreactivity in the sections were graded as none (-), mild (+), moderate (++) and severe (+++).

## Cell experiments

WT HepG2 was included in this study and cultured by Roswell Park Memorial Institute (RPMI, R2405, Sigma Aldrich) 1640 Medium with 10% fetal bovine serum (FBS) and 1% penicillin – streptomycin (PS). To induce a steatosis model into WT HepG2 cells, high glucose Dulbecco’s Modified Eagle Media (DMEM, D0822/D5671, Sigma-Aldrich) containing 10% FBS and 1% PS was used as medium for one week, together with 10μg/mL insulin and 10μM T0901317. All of cells were incubated in the incubator with 5% CO2 and 37°C (Thermo SCIENTIFIC). Fresh medium was provided every two days.

Nine drugs were tested in this study and all of them were bought from commercial companies as shown in Table S7. All the drugs were diluted in Dimethyl sulfoxide (DMSO, 41639, Sigma-Aldrich) and stored in −20°C.

As for drug-screening, different numbers of HepG2 cells were seeded for distinct experiments. More in details, 40,000 cells and 500,000 cells were used for 2-day-drug treatment in 96-well-plate and 6-well plate, respectively, while 80,000 cells and 300,000 cells were used for one-week drug treatment in 96-well plate and 6-well plate, respectively. Detailed drug dosages could be found in Table S8. Fresh medium was provided every two days.

## Transcriptomic data from human patients

We validated the key genes we identified in mouse fed with HSD by performing differential expression analysis for the publicly available human patient datasets where the study details and its characteristics are also reported (Govaere et al., 2020).

## RNA extraction and sequencing

Total RNA was isolated from homogenized heart tissue using RNeasy Fibrous Tissue Mini Kit (Qiagen). cDNA was synthesized with the high-capacity cDNA Reverse Transcription Kit (Applied Biosystems) and random primers. mRNA expression of genes of interest was analyzed with TaqMan real-time PCR in a ViiA™ 7 system (Applied Biosystems). RNA sequencing library were prepared with Illumina RNA-Seq with Poly-A selections. Subsequently, the libraries were sequenced on NovaSeq6000 (NovaSeq Control Software 1.6.0/RNA v3.4.4) with a 2×51 setup using ‘NovaSeqXp’ workflow in ‘S1’ mode flow cell. The Bcl was converted to FastQ by bcl2fastq_v2.19.1.403 from CASAVA software suite (Sanger/phred33/Illumina 1.8+ quality scale).

## Lipid quantification

Lipids were extracted as described previously (Lofgren et al, 2012). Internal standards were added during the extraction. Lipids were analysed using a combination of HPLC and mass spectrometry as described (Stahlman et al, 2013). Briefly, straight-phase HPLC was used to purify ceramides (CER). Cholesteryl ester (CE), triacylglycerol (TAG), phosphatidylethanolamine (PE), phosphatidylcholine (PC), and sphingomyelin (SM) were quantified using a QTRAP 5500 mass spectrometer (Sciex, Concord, Canada) equipped with a robotic nanoflow ion source, TriVersa NanoMate (Advion BioSciences, Ithaca, NJ). CER were analysed using reversed- phase HPLC coupled to a triple-quadrupole Quattro Premier mass spectrometer (Waters, Milford, MA, USA).

## Western blot

Protein content levels were detected by western blot. After the respective treatment of drugs, the cells pellet was lysed in cell lytic M, followed by protein extraction from supernatant (10min, 13,000r/min centrifuge). Extracted cell lysate was stored in −20°C. Proteins were quantified through Bovine albumin serum (BAS) standard curve with protein assay dye reagent (BIO-RAD). The proteins were separated by SDS-PAGE with 100v in 90 min by mini-protein TGX precast gels (4-15%, 15 wells, 15μl/well, BIO-RAD), and then transfer to Mini PVDF transfer package through transfer system (Transfer-Turbo, BIO-RAD). After 30min blocking with 5% skim milk in cold room (4°C) and washing with TBST, membrane was cut by the size of interested proteins. Primary antibodies were added into the corresponding membrane sections for overnight binding. Then the membranes were washed with TBST, and the secondary antibodies were incubated for 1h in cold room. The following antibody were used: (Goat pAb to Rb lg, ab205718, abcam; Rb pAb to beta-actin, ab8227, abcam; anti-pklr, HPA006653, SIGMA; Rb pAb to fatty acid synthase, ab99539, abcam). The binds were imaged by immunostaining and machine (ImageQuant LAS 500). All of the antibodies were purchased from Abcam online, and β-actin was used as global normalized protein. After measurement, the strength of signal for each protein was quantified by ImageJ. The background noise was removed, and all of the results were normalized by the levels of β-actin.

## Cell viability assay & Triglyceride assay for in vitro experiment

Cell viability and lipids content were detected on the same batch of cells. After one-week treatment of drugs in steatosis model, drug cytotoxicity was determined by Cell Counting Kit 8 (CCK-8, Sigma-Aldrich), and triglyceride levels were detected by Triglyceride Quantification Assay (TAG) kit (Abcam, ab65336). Then the absorbance was measured at 450 nm and 570 nm, respectively. To normalize triglyceride content level, average values of cell viability were applied.

## Transcriptomics data

The raw RNA-sequencing results were processed using Kallisto (Bray et al., 2016) with index file generated from the Ensembl mouse reference genome (Release-92) (Zerbino et al., 2018). The output from Kallisto, both estimated count and TPM (Trancript per kilobase million), were subsequently mapped to gene using the mapping file retrieved from Ensembl BioMart website, by filtering only protein coding genes and transcripts. Genes with mean expression less than 1 TPM in each condition were filtered. For data exploration, we used PCA from sklearn package (https://dl.acm.org/doi/10.5555/1953048.2078195pi) in Python 3.7 and used TPM values as the input.

Subsequently, we performed differential gene expression analysis using DESeq2 package in R (Love et al., 2014). To define a gene as differentially expressed (DEGs), a gene has to fulfil a criterion of FDR < 5%. The results of differential expression analysis were then used for functional analysis.

We performed functional analysis using the R package PIANO (Varemo et al., 2013). As the input, we used the fold changes and p-values from the DESeq2, and also KEGG pathways gene-set collections from Enrichr. To define a process or pathway as significant, we used a cut off of FDR < 10% for the distinct direction of PIANO (both up and down).

## Identification of *Pklr* gene signature

The *Pklr* signature was determined by meta-analysis of three individual RNA-seq datasets. The first two datasets were obtained from our previous study (Liu et al., 2019) where we used siRNA/plasmid to inhibit/over-express *Pklr* in HepG2 cell line, respectively. Raw sequencing data was mapped by Kallisto based on GRCh38 (gencode v23). We performed differential express analysis with R package DESeq2 for the two datasets. The results of DEGs were recognized as gene signature, which respectively referred to *Pklr* inhibition signature and *Pklr* over-expression signature hereafter. Next, we collected RNA-seq data of liver biopsies from 110 healthy donors generated by the Genotype-Tissue Expression (GTEx) project. After removing low expressed genes (TPM<1), we calculated the spearman correlation coefficient between *Pklr* and other genes, referred to Pklr co-expression signature. In summary, we detected three *Pklr*-related gene signature, namely *Pklr* co-expression signature (N = 11733), *Pklr* inhibition signature (N = 16399) and *Pklr* over-expression signature (N = 14903).

The next step was to generate a *Pklr* consensus signature by aggregating the three-candidate gene lists. First of all, genes were defined as *Pklr* positively related genes if they met three conditions at the same time: positive log2 fold change values in *Pklr* over-expression signature group; negative log2 fold change values in Pklr inhibition signature group; positive correlation coefficient in *Pklr* co-expression signature. The definition of *Pklr* negatively correlated genes was the opposite. These *Pklr* positively/negatively correlated genes generated a list of *Pklr* consensus correlated genes (N=4994).

To rank these consensus genes by the level of relationship with *Pklr* in a uniform standard, we calculated a combined z-score for them by aggregating the P-values from the three lists of *Pklr* signature. Specifically, for each list of P-values, we transformed the P-Values (two-tailed significance test) to z-score as follow:

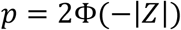

where Φ is the standard normal cumulative distribution function (Lee et al., 2016).

Then we calculated a combined z-score for 4998 combined *PKL* correlated genes. The equation is as follow:

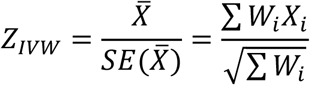

where X is the z score from three independent results and W is the weight (set as 1 for all of three results) and IVW is the Inverse variance-weighted (IVW) average method (Lee et al., 2016).

Consequently, we were able to order the 4998 Pklr consensus correlated genes by the aggregated z-scores, and these z-scores were defined as our final *Pklr* consensus signature (Table S5). Notably, z-scores of negatively correlated genes were labelled as negative values and vice versa, in order to reflect the direction of genes.

## Drug repositioning

The fifth level of LINCS L1000 phase 2 data were downloaded (GEO: GSE70138) and used for detecting the connectivity of small molecules and gene signatures. HepG2 cell line data was specifically parsed from the whole file, containing differential expressed gene signatures after small molecules’ treatment. Then the Spearman correlation coefficient was calculated between our *Pklr* consensus signatures and these drug-perturbed gene signatures, focusing on the 3698 pair-wised genes (Table S5). P-Values were adjusted based on the Benjamini-Hochberg method. Drugs were ranked by r value, and we considered top 10 drugs as potential *Pklr* inhibitors (r < −0.4, FDR < 1E-95). In total, there were 10 drugs extracted, and nine of them that we could purchase were selected for wet lab validation afterward.

## Data and code availability

All raw RNA-sequencing data generated from this study can be accessed through accession number GSE. Codes used during the analysis are available on https://github.com/sysmedicine/pklr

## Reference

Asher, G., and Schibler, U. (2011). Crosstalk between components of circadian and metabolic cycles in mammals. Cell Metab 13, 125–137.

Cerbone, A., Toaldo, C., Pizzimenti, S., Pettazzoni, P., Dianzani, C., Minelli, R., Ciamporcero, E., Roma, G., Dianzani, M.U., Canaparo, R., et al. (2012). AS601245, an Anti-Inflammatory JNK Inhibitor, and Clofibrate Have a Synergistic Effect in Inducing Cell Responses and in Affecting the Gene Expression Profile in CaCo-2 Colon Cancer Cells. PPAR Res 2012, 269751.

Charni-Natan, M., Aloni-Grinstein, R., Osher, E., and Rotter, V. (2019). Liver and Steroid Hormones-Can a Touch of p53 Make a Difference? Front Endocrinol (Lausanne) 10, 374.

Chella Krishnan, K., Floyd, R.R., Sabir, S., Jayasekera, D.W., Leon-Mimila, P.V., Jones, A.E., Cortez, A.A., Shravah, V., Peterfy, M., Stiles, L., et al. (2021). Liver Pyruvate Kinase Promotes NAFLD/NASH in Both Mice and Humans in a Sex-Specific Manner. Cell Mol Gastroenterol Hepatol 11, 389–406.

Chella Krishnan, K., Kurt, Z., Barrere-Cain, R., Sabir, S., Das, A., Floyd, R., Vergnes, L., Zhao, Y., Che, N., Charugundla, S., et al. (2018). Integration of Multi-omics Data from Mouse Diversity Panel Highlights Mitochondrial Dysfunction in Non-alcoholic Fatty Liver Disease. Cell Syst 6, 103–115 e107.

Fabbrini, E., Sullivan, S., and Klein, S. (2010). Obesity and nonalcoholic fatty liver disease: biochemical, metabolic, and clinical implications. Hepatology 51, 679–689.

Gluchowski, N.L., Becuwe, M., Walther, T.C., and Farese, R.V., Jr. (2017). Lipid droplets and liver disease: from basic biology to clinical implications. Nat Rev Gastroenterol Hepatol 14, 343–355.

Haas, J.T., Francque, S., and Staels, B. (2016). Pathophysiology and Mechanisms of Nonalcoholic Fatty Liver Disease. Annu Rev Physiol 78, 181–205.

Hansmannel, F., Mordier, S., and Iynedjian, P.B. (2006). Insulin induction of glucokinase and fatty acid synthase in hepatocytes: analysis of the roles of sterol-regulatory-element-binding protein-1c and liver X receptor. Biochem J 399, 275–283.

Hosseini, H., Teimouri, M., Shabani, M., Koushki, M., Babaei Khorzoughi, R., Namvarjah, F., Izadi, P., and Meshkani, R. (2020). Resveratrol alleviates non-alcoholic fatty liver disease through epigenetic modification of the Nrf2 signaling pathway. Int J Biochem Cell Biol 119, 105667.

Kanehisa, M., Furumichi, M., Sato, Y., Ishiguro-Watanabe, M., and Tanabe, M. (2021). KEGG: integrating viruses and cellular organisms. Nucleic Acids Res 49, D545–D551.

Lee, S., Zhang, C., Liu, Z., Klevstig, M., Mukhopadhyay, B., Bergentall, M., Cinar, R., Stahlman, M., Sikanic, N., Park, J.K., et al. (2017). Network analyses identify liver-specific targets for treating liver diseases. Mol Syst Biol 13, 938.

Liu, Z., Zhang, C., Lee, S., Kim, W., Klevstig, M., Harzandi, A.M., Sikanic, N., Arif, M., Stahlman, M., Nielsen, J., et al. (2019). Pyruvate kinase L/R is a regulator of lipid metabolism and mitochondrial function. Metab Eng 52, 263–272.

Mardinoglu, A., Uhlen, M., and Boren, J. (2018). Broad Views of Non-alcoholic Fatty Liver Disease. Cell Syst 6, 7–9.

Muller, A., and Florek, M. (2010). 5-Azacytidine/Azacitidine. Recent Results Cancer Res 184, 159–170.

Poirier, S., Samami, S., Mamarbachi, M., Demers, A., Chang, T.Y., Vance, D.E., Hatch, G.M., and Mayer, G. (2014). The epigenetic drug 5-azacytidine interferes with cholesterol and lipid metabolism. J Biol Chem 289, 18736–18751.

Stenvers, D.J., Scheer, F., Schrauwen, P., la Fleur, S.E., and Kalsbeek, A. (2019). Circadian clocks and insulin resistance. Nat Rev Endocrinol 15, 75–89.

Subramanian, A., Narayan, R., Corsello, S.M., Peck, D.D., Natoli, T.E., Lu, X., Gould, J., Davis, J.F., Tubelli, A.A., Asiedu, J.K., et al. (2017). A Next Generation Connectivity Map: L1000 Platform and the First 1,000,000 Profiles. Cell 171, 1437–1452 e1417.

Tana, C., Ballestri, S., Ricci, F., Di Vincenzo, A., Ticinesi, A., Gallina, S., Giamberardino, M.A., Cipollone, F., Sutton, R., Vettor, R., et al. (2019). Cardiovascular Risk in Non-Alcoholic Fatty Liver Disease: Mechanisms and Therapeutic Implications. Int J Environ Res Public Health 16.

Udoh, U.S., Valcin, J.A., Swain, T.M., Filiano, A.N., Gamble, K.L., Young, M.E., and Bailey, S.M. (2018). Genetic deletion of the circadian clock transcription factor BMAL1 and chronic alcohol consumption differentially alter hepatic glycogen in mice. Am J Physiol Gastrointest Liver Physiol 314, G431–G447.

Varemo, L., Nielsen, J., and Nookaew, I. (2013). Enriching the gene set analysis of genome-wide data by incorporating directionality of gene expression and combining statistical hypotheses and methods. Nucleic Acids Res 41, 4378–4391.

